# The heme exporter FLVCR regulates mitochondrial biogenesis and redox balance in the hematophagous insect *Rhodnius prolixus*

**DOI:** 10.1101/2023.08.07.552335

**Authors:** Ana Beatriz Walter-Nuno, Mabel Taracena-Agarwal, Matheus P. Oliveira, Marcus F. Oliveira, Pedro L. Oliveira, Gabriela O. Paiva-Silva

## Abstract

Heme is a prosthetic group of proteins involved in vital physiological processes in aerobic organisms. It participates in redox reactions crucial for cell metabolism due to the variable oxidation state of its central iron atom. However, excessive heme can be cytotoxic due to its prooxidant properties. Therefore, the control of intracellular heme levels ensures the survival of organisms, especially those that deal with high concentrations of heme during their lives, such as hematophagous insects. The feline leukemia virus C receptor (FLVCR) is a membrane protein responsible for heme transport in mammalian cells. In our study, we found that RpFLVCR serves as a heme exporter in the midgut of the hematophagous insect *Rhodnius prolixus*, a vector for Chagas disease. Silencing RpFLVCR decreased hemolymphatic heme levels and increased the levels of intracellular dicysteinyl-biliverdin, a product of *R. prolixus* heme degradation, indicating heme retention inside midgut cells. FLVCR silencing led to increased expression of heme oxygenase (HO), ferritin, and mitoferrin mRNAs while downregulating the iron importers Malvolio 1 and 2. In contrast, HO gene silencing increased FLVCR and Malvolio expression and downregulated ferritin, revealing crosstalk between heme degradation/export and iron transport/storage pathways. Furthermore, RpFLVCR silencing strongly increased oxidant production and lipid peroxidation, reduced cytochrome c oxidase activity and activated mitochondrial biogenesis, effects not observed in RpHO-silenced insects. These data support FLVCR function as a heme exporter, playing a pivotal role in heme/iron metabolism and maintenance of redox balance, especially in an organism adapted to face extremely high concentrations of heme.

## Introduction

Heme, an iron-protoporphyrin IX complex, plays a crucial role in the aerobic metabolism of most organisms. It is a prosthetic group of heme proteins that is involved in a vast array of cellular processes, such as antioxidant protection and xenobiotic detoxification. Due to its ability to bind diatomic gases, heme is involved in oxygen transport and NO- and CO-dependent signaling pathways. Moreover, it is directly involved in most oxidative reactions required for electron transfer and energy transduction in the mitochondria (1–3).

To safeguard cellular integrity and its function, it is essential to maintain appropriate levels of intracellular heme, as excess heme can exert cytotoxicity either by directly promoting oxidative damage to biomolecules (4–6) or by altering membrane permeability (7). In most aerobic organisms, heme homeostasis is maintained by orchestrating heme synthesis, degradation, and transport between and within cells, according to the supply and demand of this metallocofactor (8–12).

The control of heme homeostasis is particularly critical for hematophagous insects, including vectors of human diseases, which deal with large amounts of this molecule released during the digestion of the host’s blood (13). In fact, a variety of mechanisms to counteract the harmful effects of free heme and generated reactive oxygen species (ROS) have evolved in these animals (13). Some of them represent obvious adaptations of organisms that face an overload of heme, such as the formation of insoluble heme aggregates (hemozoin) (14,15), sequestration by heme-binding proteins associated with the extracellular matrix (16) and unique heme enzymatic degradation pathways that result in the production of water-soluble modified biliverdins (17,18). While the biochemical aspects of heme synthesis and degradation have already been described in some of these blood-sucking arthropods (17–20), the precise mechanisms involved in transmembrane heme transport remain largely unexplored. In ticks, an ABCB10 homolog has been implicated in heme transport (21). In contrast, in *Aedes aegypti*, exposing cells and tissues to low or high heme does not result in positive identification of heme transporters (22).

In mammals, cellular transmembrane heme export is attributed to feline leukemia virus subgroup C receptor 1 (FLVCR1). FLVCR1 was originally identified as the cell surface receptor for feline leukemia virus subgroup C (23,24), but a role in heme metabolism soon became apparent (25). Flvcr1a is one of the two isoforms codified by the Flvcr1 gene. It is a member of the SLC49 family of the major facilitator superfamily of secondary transporters, capable of transporting small solutes in response to chemiosmotic ion gradients (26,27). Flvcr1a mRNA is widely expressed in various mammalian organs, such as the bone marrow, liver, duodenum, lungs, kidneys, spleen, brain and placenta (28–30). The role of FLVCR1a (herein referred to as FLVCR) as a heme exporter has been described in different cell types, including erythroid cells, macrophages, hepatocytes, enterocytes and endothelial cells (25,28,31–33). Its contribution to the maintenance of heme homeostasis is particularly critical in erythroid cells, where heme is abundantly produced during erythropoiesis (34). In these cells, FLVCR ensures the efficient transport of excess unbound heme molecules, possibly preventing their toxic accumulation (28). However, the role of cellular heme efflux ascribed to FLVCR as an antioxidant mechanism is still controversial (35). The FLVCR1-related protein FLVCR2 has been proposed to be a heme importer in mammals (36) but also seems to be involved in calcium transport (37). Additionally, a member of the FLVCR family has been described in *Leishmania major* (LmFLVCRb) that is capable of taking up porphyrins and heme inclusively from the extracellular medium (38).

As previously described, a single FLVCR ortholog has been identified in the genome of *Rhodnius prolixus*, a vector of Chagas disease (39). RpFLVCR has the 12 transmembrane canonical domains that are typical of these proteins. RpFLVCR knockdown impacts the survival of nymphs and adults as well as oogenesis, embryogenesis and molting during their life cycle, indicating that fluctuations in the levels of this protein can be deleterious to insect development (39).

In this study, we present evidence that RpFLVCR acts as a heme transporter in *R. prolixus* and that heme efflux from the midgut lumen to the hemolymph is dependent on RpFLVCR. This process is critical for maintainingintestinal homeostasis and redox balance by modulating the expression of genes involved in iron/heme metabolism and antioxidant protection. Furthermore, RpFLVCR knockdown induces mitochondrial biogenesis with altered function, highlighting its paramount role in mitochondrial physiology in *R. prolixus*.

## Results and Discussion

### RpFLVCR acts as a heme exporter in Rhodnius prolixus

Heme in cells and extracellular fluids is rarely found as free heme and is more precisely described as “labile heme”. Labile heme is exchangeable between distinct cellular targets, including low-molecular-weight ligands or proteins, nucleic acids or phospholipid membranes (40). In vertebrates, two main mechanisms are known to be involved in the control of intracellular levels of labile heme: enzymatic degradation by heme oxygenase and heme transport provided by membrane transporters (41–43). Although heme degradation pathways have been documented in insects (17,18,44), the proteins responsible for the import and export of heme by cells remain undescribed. Heme molecules provided in the bloodmeal reach the hemolymph by crossing the midgut epithelium of the kissing bug *R. prolixus* (45), thus revealing a mechanism that provides the transfer of heme molecules from the intestinal lumen of blood-sucking insects through enterocytes to the hemolymph.

In the genome of *Rhodnius prolixus,* a single FLVCR ortholog has been identified, and gene silencing strongly affects insect reproduction, development and survival after a blood meal (39). Thus, we decided to analyze whether, as in vertebrates, this transporter is involved in the export of intracellular heme in *R. prolixus*.

Circulating levels of heme in hemolymph were measured in RpFLVCR-silenced females and used as a readout of heme export activity. Figure 1A shows that RpFLVCR-silenced females had lower heme concentrations in their hemolymph than dsMAL-injected controls. If RpFLVCR works as a heme exporter, its knockdown should lead to an increase in intracellular heme followed by heme degradation by heme oxygenase (HO). As observed in the midgut tissue, the levels of Rp-biliverdin, a breakdown product of heme catabolism (17), were higher in RpFLVCR-silenced females than in control females (Figure 1B). These results indicate that FLVCR silencing promotes an increase in the intracellular labile heme pool and support the hypothesis that RpFLVCR acts as a transmembrane exporter of heme from the intestinal epithelium to the hemolymph.

**Figure 1:**
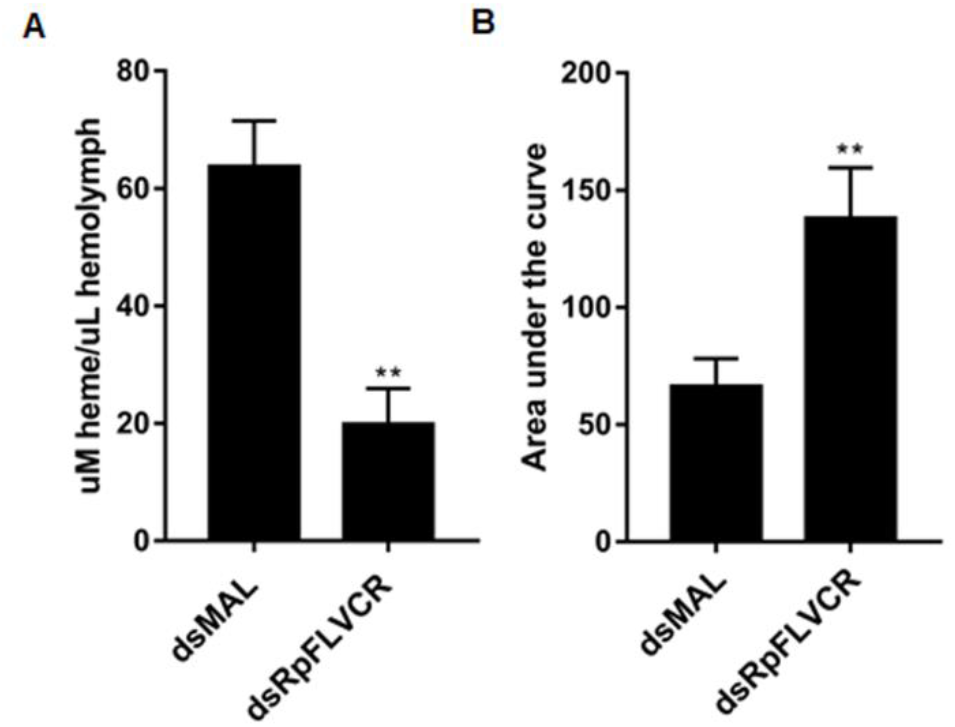
FLVCR acts as a heme exporter in *R. prolixus*. Adult females were injected with dsRpFLVCR or dsMAL (control group) and fed on blood as described. (A) Quantification of heme present in the hemolymph of females injected with dsRpFLVCR and dsMAL. The amount of heme present in hemolymph was determined by the alkaline pyridine method. The data are presented as the mean ± SE for 10 individual determinations per experiment of four independent experiments. Statistical analysis between the two groups was performed using Student’s *t* test. (B) Quantification of biliverdin present in the midgut of RpFLVCR-silenced insects. The amount of biliverdin was measured as described in Section 2 by HPLC. The data are presented as the mean ± SE for n=5 of three independent experiments. Statistical analysis between the two groups was performed using Student’s *t* test. ** *p*< 0.01

### Knockdown of RpFLVCR modulates the expression of genes involved in heme and iron metabolism

It is well established in the literature that heme is a ubiquitous molecule involved in several cellular processes, including signal transduction and transcriptional modulation (46–54). Thus, possible variations in heme levels caused by RpFLVCR KD could impact the transcription of other genes involved in iron and heme metabolism. Thus, we sought to observe phenotypes that could be associated with variations in intracellular heme levels. We analyzed the mRNA levels of genes encoding key proteins such as HO (heme degradation), Malvolium 1 and 2 (NRAMP orthologs from mammals involved in iron import), mitoferrin (Mfrn, a mitochondrial iron importer) and ferritin (Fer, an intracellular iron-storage protein). RpFLVCR knockdown promoted increases in the expression of HO, Mfrn, and Fer and decreases in the expression of the iron importers MVL1 and MVL2 (Figure 2). These results suggest that the increase in HO expression may have been a compensatory mechanism to control intracellular heme levels that were not exported due to RpFLVCR silencing. Increased expression of intracellular iron-binding proteins, such as ferritin (Fer) and mitoferrin (Mfrn), was required to prevent oxidative damage caused by iron from heme porphyrin ring breakage. On the other hand, with the increased availability of intracellular iron, import by MLV proteins became unnecessary, explaining the decrease in MLV expression (Figure 2B). Interestingly, HO KD caused an opposite effect on gene expression in comparison to that of RpFLVCR silencing, leading to increased expression of the RpFLVCR, MVL1 and MVL2 genes and decreased mRNA levels of ferritin (Figure 2C and 2D). These results suggested that the reduction in HO expression increased intracellular heme but decreased labile iron levels. These results are consistent with increased RpFLVCR transcription working to remove excess heme. A decrease in free iron availability would promote MVL importer expression and, conversely, lead to reductions in the expression of iron-chelating genes, such as ferritin and mitoferrin.

**Figure 2:**
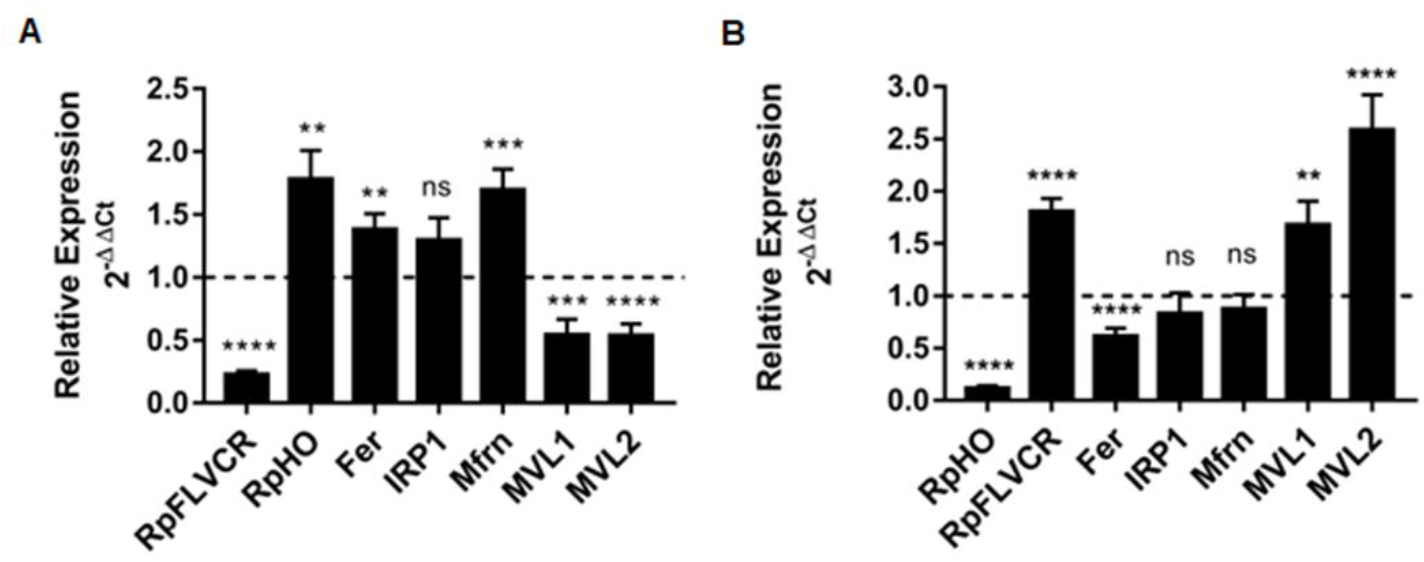
FLVCR silencing is able to modulate genes from heme and iron metabolism. Starved adult females were injected with dsRpRFLVCR (A) or dsRpHO (B). Insects were fed on blood 48 h after injection. Total RNA was extracted from the foregut and hindgut, and the mRNA levels of genes involved in iron and heme metabolism were analyzed by quantitative RT‒PCR. The EF-1 gene was used as an endogenous control. The result was normalized to animals injected with dsMAL (dashed line). The data are presented as the mean ± SE (n= 14-15) of three independent experiments. Statistical analyses were performed using Student’s t test (experimental versus their respective controls, dsMal). *p<0.05; ** p<0.01; *** p<0.001; ****p<0.0001. RpFLVCR, feline leukemia virus C receptor FLVCR; RpHO, heme oxygenase; Fer, ferritin; IRP1, iron regulatory protein 1; Mfrn, mitoferrin; MVL1, Malvolio 1; MVL2, Malvolio 2

### RpFLVCR depletion caused redox imbalance

Heme is known to be a pro-oxidant molecule due to its ability to promote the formation of radical species (ROS). It can promptly react with organic hydroperoxides, producing alkoxyl or peroxyl lipid radicals, thus increasing lipid peroxidation (6). Indeed, the midguts of RpFLVCR-silenced insects showed increased fluorescence of the redox-sensitive probe DHE (Figure 3A), suggesting redox imbalance. This conclusion was further supported by TBARS assay analysis of lipid peroxidation levels in the midgut, which showed increased lipid peroxidation in dsRpFLVCR-injected females (Figure 3C). The impact on the midgut redox balance was also monitored by quantification of mRNA levels of antioxidant genes whose expression is usually induced in response to high levels of ROS, such as catalase (CAT), glutathione peroxidase (PHGPx), thioredoxin reductase (TrxR) and peroxiredoxins (Prx). Corroborating the DHE fluorescence and TBARS data, females with RpFLVCR gene silencing showed higher antioxidant enzyme mRNA levels than control insects (Figure 3D). Surprisingly, HO KD had no effect on the cellular redox status (Figure 3B, C and D). This result could be explained by one of two mechanisms: i) Rp FLVCR mediated export of the remaining labile heme left undegraded upon HO KD or ii) the oxidative insult to the intestinal cells upon RpFLVCR KD was derived more from iron produced via heme cleavage by HO than from an increase in the level of labile heme itself. In fact, it has already been shown that knockdown of ferritin, the main intracellular iron chelator, causes a dramatic increase in oxidant production in the *R. prolixus* midgut (39). Choosing between these alternative hypotheses is a central point for future research on redox metabolism in blood-sucking insects.

**Figure 3.**
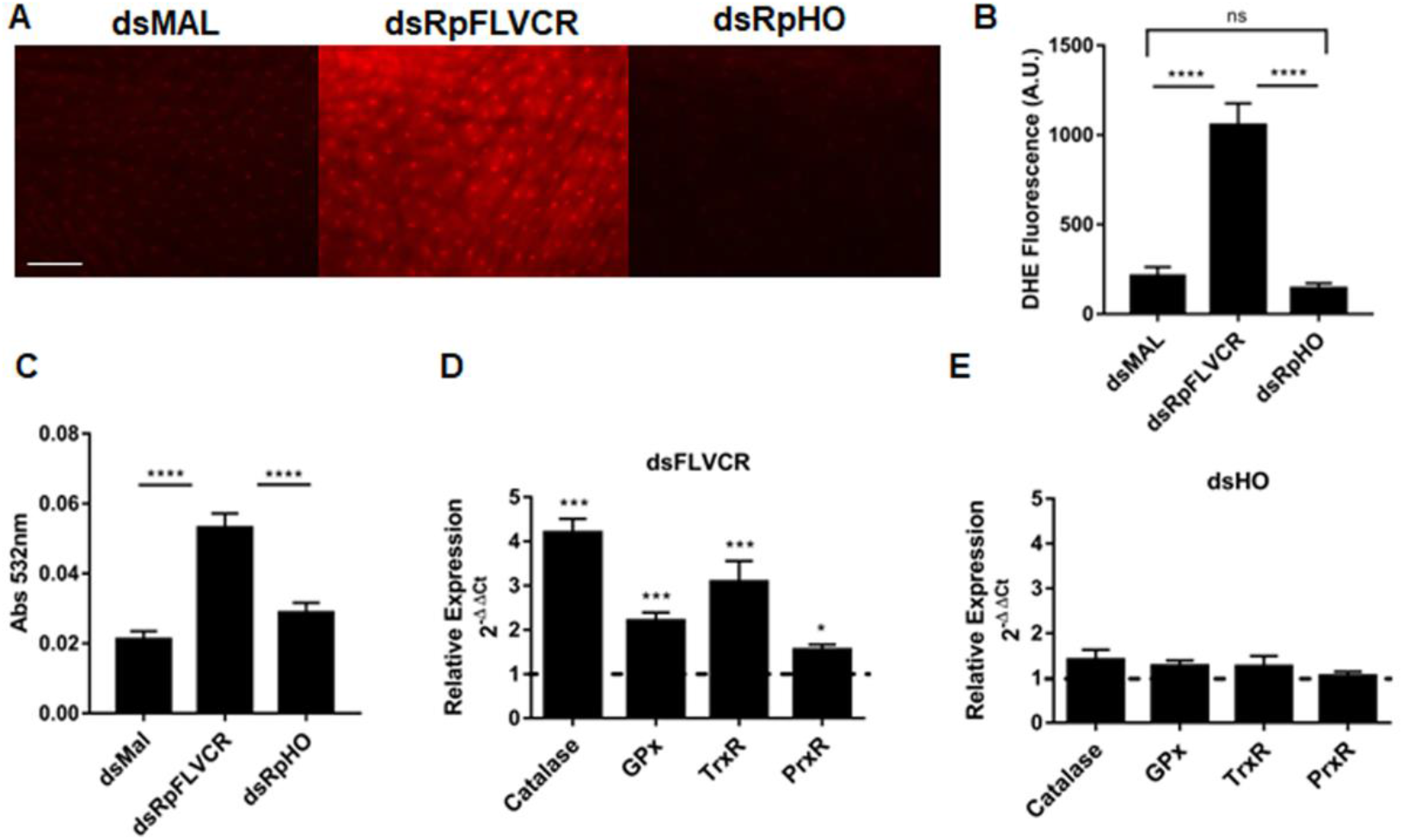
FLVCR silencing increases ROS levels in midgut cells. (A) ROS levels were evaluated from oxidized DHE fluorescence in cells from blood-fed females previously injected with dsRpFLVCR, dsRpHO or dsMal (control). Representative images of insects for each experimental condition are shown, scale bar 2 μm. Quantitative analysis of DHE fluorescence in 10 individual intestines from three independent experiments. The data shown are the mean ± SE, **** p<0.0001 for one-way ANOVA with Tukey’s posttest. (B) Lipid peroxidation of hemolymph was quantified by the TBARS assay as described. The data shown are the mean ± SE (n= 4) from three independent experiments. **** p<0.0001 for one-way ANOVA with Tukey’s posttest. (C-D). The expression of selected antioxidant genes was evaluated by quantitative RT‒PCR in insects with silenced RpFLVCR (C) or RpHO (D). The result was normalized to that of animals injected with dsMAL (dashed line). Statistical analyses were performed using Student’s t test (experimental versus their respective controls, dsMal). The data shown are the mean ± SE, *p<0.05; *** p<0.001 (n=12-14).

### RpFLVCR silencing induces mitochondrial biogenesis with altered function

Heme proteins represent a significant fraction of the electron transport system (ETS) components, implying that mitochondria are the main sites of iron and heme storage in eukaryotic cells (55). Therefore, changes in heme levels in animals with RpFLVCR silencing should interfere with energy metabolism and mitochondrial function. To test this hypothesis, we first determined mitochondrial content in the intestinal cells of the silenced animals through MitoTracker Green fluorescence microscopy, as this probe accumulates preferably in mitochondria. We observed that 5 days after blood feeding, the females with silencing of the RpFLVCR gene showed a higher fluorescence than the control insects or insects with silencing of HO, suggesting an increase in the mitochondrial content (Figure 4A and 4B). This result was corroborated by the increased activity of citrate synthase, an important tricarboxylic acid cycle enzyme classically used as an index of mitochondrial content (56), in the midguts of RpFLVCR-silenced insects (Figure 4C). Surprisingly, cytochrome c oxidase activity was strongly reduced specifically in RpFLVCR-silenced insects, suggesting that *de novo* mitochondrial biogenesis generates organelles with lower electron flux and respiratory rates (Figure 4D). To gain further insights into the mechanisms underlying FLVCR KD-induced mitochondrial expansion, we measured the expression of genes that have a central role in mitochondrial biogenesis. Mitochondrial transcription factor A (TFAM) is a nuclear DNA-encoded protein responsible for the transcription and translation of mitochondrial DNA (mtDNA) (57,58). Delg (CG6338; *Drosophila* Ets-like gene) is a close homolog of mammalian NRF-2α that is critical for adjusting mitochondrial abundance (59), and its expression is frequently associated with mitochondrial biogenesis. The quantification of *R. prolixus* orthologs of these genes revealed that RpFLVCR KD promoted increases in the mRNA levels of both genes, whereas HO KD did not (Figure 4 E and F), suggesting that RpFLVCR silencing increases mitochondrial mass in the midgut in association with increased TFAM and DELG expression.

**Figure 4:**
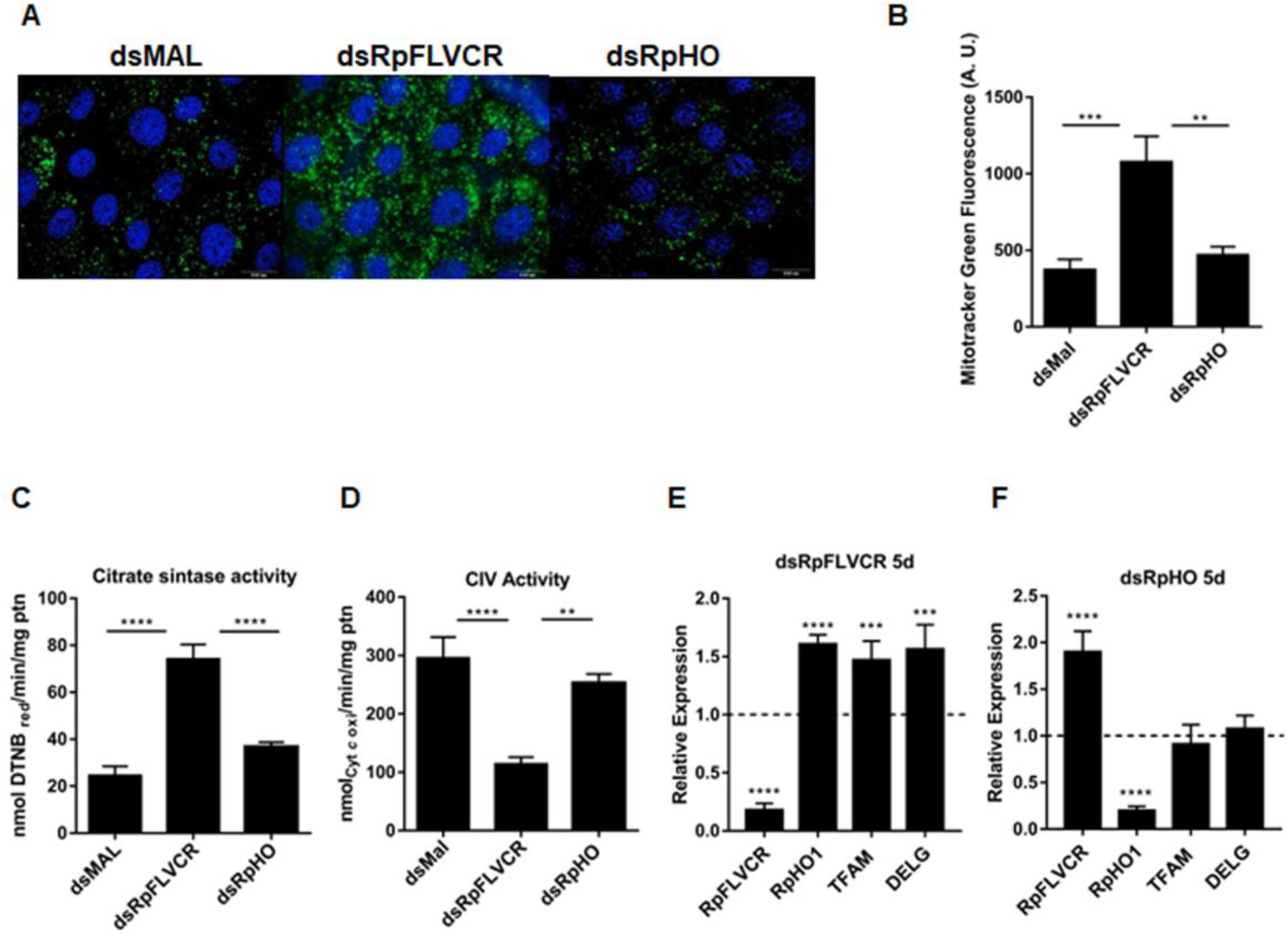
Knockdown of FLVCR increases mitochondrial biogenesis in the midgut. (A) Unfed *R. prolixus* females were injected with dsMal, dsRpFLVCR or dsRpHO; 3 days after injection, they were fed on blood. Five days after a blood meal, the insects were incubated with MitoTracker Green and DAPI as described in the Experimental section. The scale represents 20 μm. All pictures were taken with an exposure time of 260 ms. The images are representative of a total of 18 intestines analyzed. (B) Quantification of MitoGreen fluorescence using Olympus quantification software. **P≤ 0.01; *** P≤ 0.001 one-way ANOVA with Tukeýs posttest. (C) The mitochondrial content of the midgut was also determined by measuring the activity of citrate synthase in silenced females as described in the methods section. The data shown are the mean ± SE. ****P<0.0001. One-way ANOVA with Tukey’s posttest (n=7). (D) The activity of cytochrome *c* oxidase was measured using a Shimadzu spectrophotometer UV‒visible 2450. The data shown are the mean ± SE. ** P≤ 0.01; ****P≤ 0.0001, one-way ANOVA with Tukey’s posttest (n=8). (E) dsRpFLVCR- and (F) dsRpHO-injected females had their midgut dissected 5 days after feeding. The transcript levels of genes that control mitochondrial mass were measured by quantitative RT‒PCR. The result was normalized to that of animals injected with dsMAL (dashed line). Statistical analyses were performed using Student’s t test (experimental versus their respective controls, dsMal). The data shown are the mean ± SE. *** P≤ 0,001; ****P≤ 0.0001. (n= 8-11) TFAM; mitochondrial transcription factor A; DELG; CG6338/<u>D</u>rosophila <u>E</u>ts-like <u>g</u>ene.

Consistent with the finding that RpFLVCR KD promotes mitochondrial biogenesis and the knowledge that mitochondria are sources of ROS in eukaryotic cells (60–62), we noticed that the RpFLVCR KD-induced increase in DHE fluorescence upon RpFLVCR KD was strongly reduced by MitoTempo (Figure 5A). Conceivably, increased cellular oxidant production upon RpFLVCR silencing results from reduced cytochrome c oxidase activity, facilitating mitochondrial electron leakage and superoxide production. In this sense, mitochondria represent the main sites of cellular oxidant production (63), and altered mitochondrial electron flux promoted by RpFLVCR silencing (Figures 4 and 5) might contribute to superoxide production. Altogether, these results indicate that the redox imbalance observed in RpFLVCR-silenced insects is derived from mitochondrial oxidant production. However, the mechanism by which RpFLVCR KD modulates mitochondrial biogenesis and the relationship of this phenomenon with the increase in cellular oxidant levels still need to be clarified.

**Figure 5:**
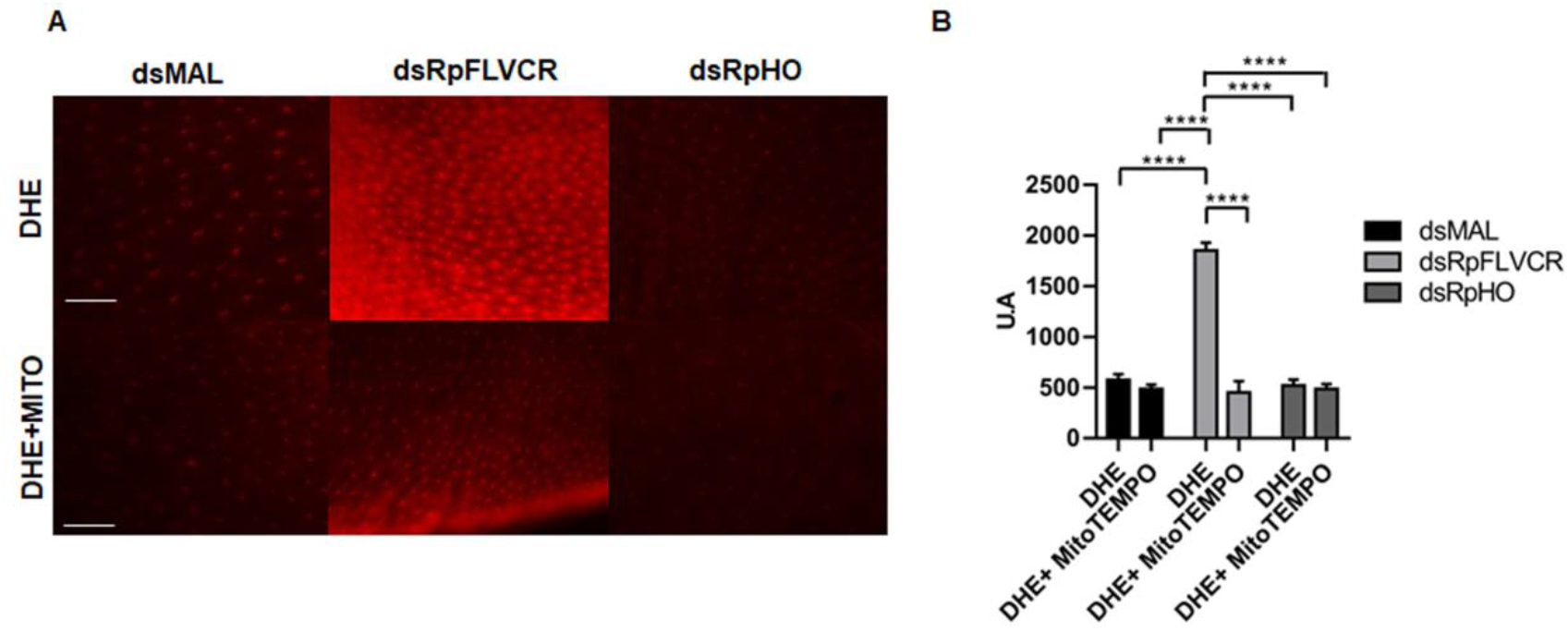
Knockdown of FLVCR promotes an imbalance in the mitochondrial redox state. (A) Anterior midguts (AMs) from FLVCR and HO-silenced females were incubated with a 50 µM solution of oxidant-sensitive fluorophore DHE or the tissues were preincubated with 50 µM MitoTEMPO, a selective mitochondrial antioxidant, prior to DHE incubation. dsMal-injected females were used as controls. DHE oxidation in the AMs was analyzed by fluorescence microscopy (Zeiss Axio Observer Microscope) using a 20× lens with an 80 ms exposure time for all conditions. The images are representative of a total of 10 midguts analyzed. Scale bar 2 µm. (B) Graph represents the quantification of fluorescence using Zeiss Axio Observer Quantification software. The data shown are the mean ± SE. ****P ≤ 0.0001 (one-way ANOVA followed by Tukey’s test).

Recently, it was shown in two elegant reports that the mammalian FLVCR works as a choline transporter instead of a heme transporter (64,65). Evidence excluding a role for this membrane protein in heme transport came from experiments showing that FLVCR KO in cells does not impact the expression of HO or other heme- and iron-related genes (64). The same report also observed alterations in mitochondria comparable to those found here but instead attributed this phenotype to deficient phospholipid biosynthesis. Here, we did not address a role of RpFLVCR in choline transport, an interesting possibility that could well explain part of the deleterious effects of RpFLVCR on insect oogenesis and embryo and nymph development (39), all biological phenomena that need large amounts of phosphatidylcholine. However, the results presented here reveal a robust impact of FLVCR on heme and iron cell biology that is unlikely to be explained by choline deficiency alone. Therefore, it is possible that FLVCR is a moonlighting transporter, with choline transport being the function for most cell types that live in the presence of heme concentrations in the low micromolar range (used in the experiments of Kenny et al. (64)), while cells that physiologically face higher heme concentrations may use FLVCR as a low-affinity but functional transporter. Vertebrate blood has approximately 150 mg/ml hemoglobin, which, upon complete digestion, should result in ∼5 mM heme. This considerable number suggests that RpFLVCR may well be such a moonlighting heme transporter. As hematophagy has arisen independently multiple times during the evolution of insects (66), further investigation with other hematophagous insect species is needed to determine how frequently the recruitment of FLVCR homologs has been a key player in the adaptation of the insect to hematophagy, as seems to be the case for *Rhodnius prolixus* intestinal epithelia.

## Experimental procedures

### Experimental Insects

Insects were taken from a colony of *R. prolixus* maintained at 28 °C and 70– 90% relative humidity under a photoperiod of 12 h of light/12 h of dark. All animals used in this work were mated females fed on rabbit blood at 3-week intervals. *R. prolixus* females injected with dsRNA were kept in individual vials maintained under the same conditions.

### Ethics Statement

All animal care and experimental protocols were conducted following the guidelines of the institutional care and use committee (Committee for Evaluation of Animal Use for Research from the Federal University of Rio de Janeiro, CAUAP-UFRJ), which are based on the National Institutes of Health Guide for the Care and Use of Laboratory Animals (ISBN 0-309-05377-3). The protocols were approved by the Committee for Evaluation of Animal Use for Research from the Federal University of Rio de Janeiro under registry number IBQM0123/22. Dedicated technicians in the animal facility at the Instituto de Bioquímica Médica Leopoldo de Meis (UFRJ) carried out all protocols related to rabbit husbandry under strict guidelines to ensure careful and consistent animal handling.

### Tissue Isolation and RNA Extraction

Posterior midguts from blood-fed females were dissected on different days after a blood meal. Total RNA was extracted from individual tissues or pools of 3–5 midguts using TRIzol reagent (Invitrogen, Carlsbad, CA, United States) according to the manufacturer’s instructions. Complementary DNA was synthesized using a High-Capacity cDNA Reverse Transcription Kit (Applied Biosystems, Foster City, CA, United States).

### dsRNA Synthesis and Gene Silencing Assays

Fragments of 320–400 bp were amplified by PCR using cDNA from midgut epithelia from blood-fed females (48 h after feeding) produced as described above. The following conditions were used for amplification: one cycle for 5 min at 95 °C followed by 40 cycles of 30 s at 95 °C, 30 s at 63 °C, and 1 min at 72 °C, with a final step of 10 min at 72 °C. The oligonucleotides used for amplification of templates for dsRNA synthesis are listed in Supporting Table S1. Amplified cDNAs were used as a template for dsRNA synthesis using a MEGAScript RNAi Kit (Ambion Inc., Austin, TX, United States) according to the manufacturer’s instructions. The maltose-binding protein (MAL) gene from *Escherichia coli* (gene identifier 7129408) in a pBlueScript KS (Stratagene) was amplified by PCR using T7 minimal promoter primers under the following conditions: one cycle for 10 min at 95 °C followed by 40 cycles of 15 s at 95 °C, 15 s at 45 °C, and 45 s at 72 °C, with a final step of 10 min at 72 °C. The PCR product produced was used as a template for MAL dsRNA synthesis and as a control in the gene silencing assays. Following *in vitro* synthesis, all dsRNAs were purified according to the manufacturer’s instructions. For gene silencing, 1 μg of dsRNA specific for RpFLVCR (dsRpFLVCR), heme oxygenase (dsHO) or MAL (dsMAL) was injected into the starved insect hemocoel. Insects were fed on rabbit blood 48 h after injection. At different times after feeding, midguts were dissected, total RNA was extracted as described above, and gene silencing efficiency was evaluated by quantitative RT‒PCR. The efficiency of RpFLVCR and RpHO silencing is shown in Supporting Figure S1.

### Quantitative RT‒PCR Analysis

Quantitative gene amplification (qPCR) was performed with a StepOnePlus real-time PCR system (Applied Biosystems, Foster City, CA, United States) using Power SYBR Green PCR Master Mix (Applied Biosystems, Foster City, CA, United States) under the following conditions: one cycle for 10 min at 95 °C followed by forty cycles of 15 s at 95 °C and 45 s at 60 °C. PCR amplification was performed using the oligonucleotides specified in Supporting Table S2. *R. prolixus* elongation factor 1 gene (RPRC007684) expression was used as an internal control for normalization (67). The relative expression values based on 2^-^ΔΔCT were used in silencing gene analyses (68). Differences were considered significant when *p* < 0.05. The relative expression values based on the 2^-ΔΔ*C*^_T_ method were used only for figure construction.

### Quantification of heme content in the hemolymph

Hemolymph was collected 3 days after blood feeding from previously silenced females in tubes containing a few crystals of phenylthiourea by cutting a leg and applying gentle pressure to the insect abdomen. Qualitative analyses of heme levels were performed by diluting a hemolymph aliquot of 10 μL in 490 μL of PBS pH 7.4, and the light absorption spectra were analyzed between 300 nm and 800 nm in a Shimadzu UV-2550 spectrophotometer. Absolute quantification of total heme was determined by the alkaline pyridine-hemochrome method using the reduced minus oxidized spectra as described elsewhere (69).

### Quantification of biliverdin in the midgut

Two days prior to the blood meal, adult females were injected with 1 μL of dsRNA (1 μg/μL) into the thoracic cavity. Five individual midguts were dissected 3 days after feeding; individually homogenized with 200 μL of 5% acetonitrile and 0.05% TFA, pH 2.0, (used as solvent for the HPLC column); and centrifuged for 15 min at 12,000×g. The supernatant was applied onto a Shimadzu CLC-ODS C18 column (15 mm × 22 cm) equilibrated with the same solvent using a flow rate of 0.4 mL/min using an LC-10AT device (Shimadzu, Tokyo, Japan) equipped with a diode array detector (SPDM10A). After 10 min, an acetonitrile linear gradient (5–80%) was applied for 10 min followed by 20 min of 80% acetonitrile with 0.05% TFA, pH 2.0. The biliverdin peak was identified and quantified as previously described (17).

### Analysis of oxidant status in different tissues

Lipid peroxidation levels in the hemolymph of silenced females were assessed 3 days after a blood meal by quantifying the levels of thiobarbituric acid reactive substances (TBARS) at 532 nm in a Molecular Devices Spectra Max M5 spectrophotometer as described elsewhere (70). Additionally, to evaluate oxidant levels in the midgut, starved females were injected with 1 μg of RpFLVCR dsRNA, HO dsRNA or MAL dsRNA (control). Two days after injection, insects were fed with rabbit blood, and their wings, legs and dorsal cuticle were removed by dissection. The insect hemocoel was filled with 50 μM of the oxidant-sensitive fluorophore dihydroethidium (hydroethidine; DHE; Invitrogen, Carlsbad, CA, United States) diluted in Leibovitz-15 medium containing 5% fetal bovine serum or with DHE and 5 µM MitoTEMPO (Santa Cruz, Dallas, TX, United States). The midguts were incubated in the dark at 28 °C for 20 min, washed with 0.15 M NaCl, and immediately transferred to a glass slide for fluorescence microscopy analysis, as previously described (71). Quantitative evaluation of fluorescence levels was performed by acquiring images under identical conditions using an objective of 20X and 90 ms exposure time in each experiment. The images were acquired in a Zeiss Observer. Z1 with a Zeiss Axio Cam MrM Zeiss, and the data were analyzed using AxioVision version 4.8 software. The filter set (excitation BP 546/12 nm; beam splitter FT 580 nm; emission LP 590 nm) was used for DHE labeling.

### Analysis of mitochondrial content

Mitochondrial content was determined by measuring the activity of citrate synthase (CS) and by MitoTracker Green fluorescence. CS activity was assayed according to the method described by Hansen and Sidell (1983)(72), with modifications. Pools of two midguts were homogenized in 200 μL of saline solution. After 2 min of decantation, 40 μL of supernatant was incubated with 7.5 mM Tris buffer (pH 8.0) containing 50 μM DTNB (5,5′-dithiobis-(2-nitrobenzoic acid)) (Sigma‒Aldrich, MI, United States), 300 μM acetyl-CoA and 1 mM oxaloacetate. Immediately, DTNB reduction was measured for 10 min at 412 nm. The specific activity was calculated using the reduced DTNB molar extinction coefficient (**ɛ =**13.6 mM^−1^·cm^−1^). MitoTracker Green is a mitochondrial selective probe that binds covalently to mitochondrial proteins by reacting with cysteine residues and accumulates in the mitochondrial matrix (73–75). This reaction is independent of mitochondrial membrane potential and is widely used in the literature to represent mitochondrial mass (56,76). Starved adult females were injected with 1 μL of dsRpFLVCR, dsHO or dsMAL into the thoracic cavity. Two days after injection, the insects were fed on blood. Mitochondria were stained with 100 µM MitoTracker Green FM (Invitrogen, Carlsbad, CA, United States) diluted in Lebovitz-15 medium containing 5% fetal bovine serum for 30 minutes and with DAPI (0.1 mg/mL) for 10 minutes at 28 °C in the dark 5 days after feeding. Afterward, the midguts were washed with 0.15 M NaCl and immediately transferred to a glass slide for fluorescence microscopy analysis. Images were acquired in an Olympus IX81 microscope and a CellR MT20E Imaging Station equipped with an IX2-UCB controller and an ORCAR2 C10600 CCD camera (Hammamatsu). Image processing was performed with Xcellence RT version 1.2 Software.

### Cytochrome c Oxidase Activity

The activity of cytochrome *c* oxidase was measured in a total reaction volume of 1 mL using a Shimadzu UV‒visible 2450 spectrophotometer (Shimadzu Scientific Instruments, Tokyo, Japan). Enzyme activity was measured by following the decrease in absorbance caused by the oxidation of ferrocytochrome c (ɛ = 18.5 mM^−1^ cm^−1^) (77) modified according to Gaviraghi (78). The reaction mixture consisted of 100 mM potassium phosphate (pH 7.4) and 50 mM reduced cytochrome c. The reaction was initiated by the addition of freeze‒thawed mitochondria (80 μg of protein), and the reduction in absorbance at 550 nm was monitored. KCN (1 mM) was added to inhibit cytochrome *c* oxidase activity, and the resulting rate was considered the cyanide-sensitive rate of cytochrome *c* oxidation. The data are expressed as nmol of reduced cytochrome *c*/min/mg protein of isolated mitochondria.

### Statistical analysis

All analyses were performed with the GraphPad Prism statistical software package (Prism version 6.0, GraphPad Software, Inc., La Jolla, CA). Significant differences are indicated by asterisks (*, p < 0.05; **, p <0.01; ***, p < 0.001; ns, not significant), and the type of test used in each analysis is described in its respective figure legend.

## Supporting information

This article contains supporting information.

## Author contributions

ABWN, MFO and GOPS conceived and designed the experiments; ABWN, MTA and MPO performed the experiments; ABW-N, MFO, PLO and GOPS analyzed the data; ABWN wrote the original draft; all authors reviewed and edited the manuscript; and GOPS supervised the study.

## Acknowledgments

We thank all members of the Laboratório de Bioquímica de Artrópodos Hematófagos for critical suggestions and S. R. Cássia for providing technical assistance.

