## Supporting information for "The heme exporter FLVCR regulates mitochondrial biogenesis and redox balance in the hematophagous insect *Rhodnius prolixus*"

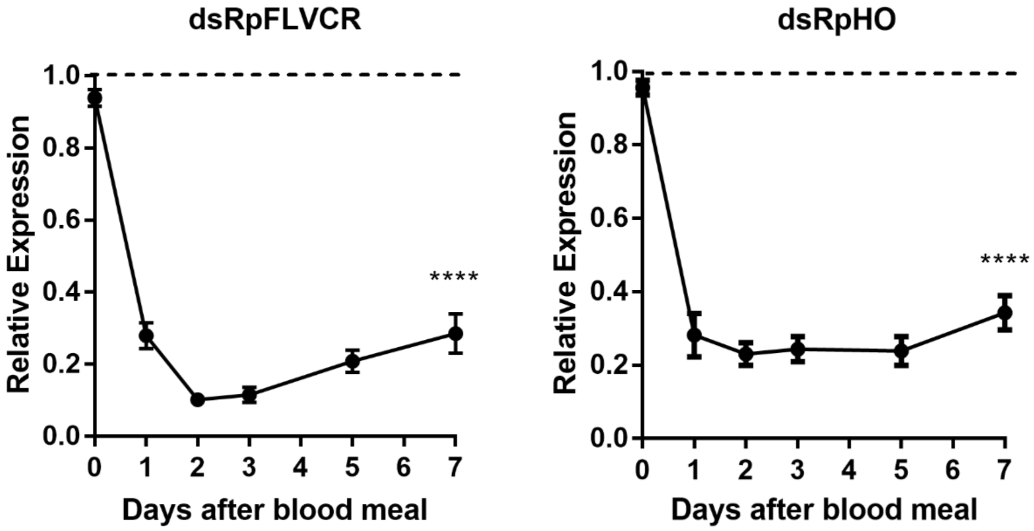


**Figure S1: dsRNA-mediated knockdown of RpFLVCR and RpHO on days after feeding**. Adult females were injected with 1 µg of dsRNA for RpFLVCR and RpHO. Insects were fed on blood 48 hours after dsRNA injection. The females were dissected, and total RNA was extracted from the posterior midgut on different days after feeding. The expression levels of the genes were determined by real-time PCR. The elongation factor 1 (EF-1) gene was used as an endogenous control. The result was normalized in relation to the animals injected with dsMAL (dashed line). The data on the graph are the mean ± SE of three independent biological replicates (pool of 3-5 midguts), each analyzed in triplicate. A two-way ANOVA with Bonferroni posttest was used to evaluate differences between the expression of the different genes and the control group at different times (**** $P<0.0001$).

**Table S1: Oligonucleotide sequences used for amplification of dsRNA synthesis templates.** Oligonucleotides were designed using Primer 3 version 0.4.0 (http://bioinfo.ut.ee/primer3-0.4.0). The T7 promoter sequence necessary for transcription is underlined. The gene sequences and accession numbers used are as deposited in the Vectorbase database (https://www.vectorbase.org/).

| Name | Accession number | Sequence |
|---|---|---|
| T7RpFLVCRFw | RPRC015407 | TAATACGACTCACTATAGGGGGTTCTGTTGTTTGTGGTTT |
| T7RpFLVCRRv | RPRC015407 | TAATACGACTCACTATAGGGTTGAGCTTCCTGCCTTCTGT |
| T7RpHOFw | RPRC006832 | TAATACGACTCACTATAGGGTGGAACAAGCTATGTCTGCAAAT |
| T7RpHORv | RPRC006832 | TAATACGACTCACTATAGGGTCTTCGTCAAGTGAATCAGCGA |

**Table S2: Oligonucleotide sequences used for real-time PCR assays.** Oligonucleotides were designed using Primer 3 version 0.4.0 (http://bioinfo.ut.ee/primer3-0.4.0). The gene sequences and accession numbers used are as deposited in the Vectorbase database (https://www.vectorbase.org/).

| Name | Accession numbers | Sequence |
|---|---|---|
| qRpFLVCRFw | RPRC015407 | CGGCTCATGACAAATCCAGG |
| qRpFLVCRRv | RPRC015407 | AATCCAATCCTCCCTGCGTC |
| qRPHOFw | RPRC006832 | CCAGAGACATTCATTCTGCATTTTAG |
| qRPHORv | RPRC006832 | GTCCTCTCCATGCCATTCAC |
| qRpEF-1Fw | RPRC007684 | GATTCCACTGAACCGCCTTA |
| qRpEF-1Rv | RPRC007684 | GCCGGGTTATATCCGATTTT |
| qRpIRP-1Fw | RPRC001246 | ACGACAGTTAGGAGTGGTAGGT |
| qRpIRP-1Rv | RPRC001246 | AGAAACCAACAGTGGCACCAT |
| qRpMfrnFw | RPRC002819 | GGAAGGCGTCGTTGCATTTT |
| qRpMfrnRv | RPRC002819 | CCACTTACCACATGCGCTTT |
| qRpMVL1Fw | RPRC006515 | AAGACCGTTCAATATCCG |

| | | |
|---|---|---|
| qRpMVL1Rv | RPRC006515 | ATGGAATAACGCAGGACC |
| qRpMVL2Fw | RPRC012012 | CATGGTGGCAGATTGTGGGC |
| qRpMVL2Rv | RPRC012012 | CCTTACAAGCGCATGAGT |
| qRpFerFw | RPRC009256 | AAAGAATGCGAAGGTTCACG |
| qRpFerFw | RPRC009256 | GCTTTTCCAGCTAAATCCCTCTG |
| qRpCATFw | RPRC007907 | TTCATCCACACGCAGAAGAG |
| qRpCATRv | RPRC007907 | GCAAGTTTCACCTCGGTCAT |
| qRpPHGPxFw | RPRC015008 | TATGGGTTTTGTTGCAGACG |
| qRpPHGPxRv | RPRC015008 | CACATTGCGATGCTACGTTC |
| qRpTrxFw | RPRC015195 | CTCGCGGTCACTCTCGGCAT |
| qRpTrxRv | RPRC015195 | CAGGTCCAGAAGCAGCGCCT |
| qRpPrxFw | RPRC015388 | TCCTGCTGATTTCACACCTG |
| qRpPrxRv | RPRC015388 | CGATCCTCATCGGAAATGAT |
| qRpTFAMFw | RPRC13701 | GCGTAGCTACAGCAAGGTTT |
| qRpTFAMRv | RPRC13701 | ACCGGCTTCCTTGGTTTAGG |
| qRpDELGFw | RPRC012582 | GAAAGCGTTGTGAGATGGGC |
| qRpDELGRv | RPRC012582 | ATGGGCAACATCCCACTGTT |

1
